# Cortical Microstructural Variations Correlate with Individual Differences in Gamified Exploration–Exploitation Behaviours

**DOI:** 10.1101/2025.10.08.681181

**Authors:** Ashley Tyrer, Niia Nikolova, Magda Dubois, Leah Banellis, Melina Vejlø, Tobias U. Hauser, Micah Allen

## Abstract

The exploration-exploitation trade-off is ubiquitous in our everyday lives, and individuals display considerable variability in their preferred decision-making strategies. Most previous work pertaining to neural signatures of exploration is restricted to functional pathways. However, the specific contributions of cortical microarchitectures to high-level cognitive processes such as decision-making are as yet unknown. Here, we investigated the neuroanatomical foundations of inter-individual variability in decision-making strategies. To this end, 122 healthy participants completed a gamified multi-armed bandit paradigm aimed at teasing apart distinct exploration-exploitation decision strategies. We also collected whole-brain quantitative MRI maps indexing microstructural features of cortical myelination and iron content. Through computational modelling, we disentangled individual-specific exploration strategies, including value-free random exploration. Whole-brain regression analyses identified significant associations between value-free exploration and increased cortical myelination in right frontal brain areas with reported links to impulsivity. By elucidating the brain microstructural correlates of distinct exploration-exploitation strategies, we aimed to further our understanding of why individuals differ in their decision-making capabilities, and how decision-making may become aberrant in mental health conditions.

## Introduction

It has been long established that individuals demonstrate considerable variability in the ways they make decisions and process information pertaining to rewards (Gershman, 2018; Schulz & Gershman, 2019). However, the neurobiological basis of these differences is not currently well characterised. A plausible explanation may be that inter-individual differences in cortical microstructures are driving this behavioural variability (Ziegler et al., 2019). Cortical microstructures such as myeloarchitecture play a vital role in the efficiency of neural signalling, by improving conduction velocity and reducing signal-to-noise ratio (Nave, 2010; Purves et al., 2001). By investigating how individual-specific differences in the brain’s structural features relate to decision-making, we can gain insight into the latent causes of such wide-ranging behavioural differences. Through this approach, we may reveal why aberrant or suboptimal decision-making strategies are often observed in individuals suffering from mental health conditions such as ADHD or depression (Must et al., 2013; Schulze et al., 2021).

Inter-individual differences in learning and decision-making become apparent when examining decision-making dilemmas such as the exploration-exploitation trade-off. In such situations, one must choose either the option with the greatest known value (exploitation), or an alternative, lesser-known option which may potentially yield yet greater rewards but at the risk of disappointing outcomes (exploration). The methods by which agents select different decision-making strategies is highly idiosyncratic and context-specific, and individuals vary greatly in their preferred decision strategies (Frank et al., 2009; Somerville et al., 2017; von Helversen et al., 2018).

Recent advances have led to the development of computational models that allow for the estimation of subject-specific parameters to describe key signatures of these exploration-exploitation behaviours (Gershman, 2018; Schwartenbeck et al., 2019). Previous work has identified the employment of exploration strategies that consider all available choices to be equally likely by disregarding all existing knowledge of the decision space, including any uncertainty and expectation estimates, thus reducing computational complexity (Daw et al., 2006; Wilson et al., 2014). This strategy is referred to as *value-free random exploration*. A recent study by Dubois et al. (2021) found clear evidence that humans employ a combination of complex resource-heavy approaches and simpler heuristic exploration strategies such as value-free random exploration, as well as uncertainty-led exploration and novelty exploration, to estimate an optimal decision method (Dubois et al., 2021). Recent work has also highlighted the neurochemical basis for decision-making, demonstrating that noradrenergic and dopaminergic activity play crucial roles in modulating exploration-exploitation strategy selection (Chakroun et al., 2019; Cremer et al., 2023; Dubois & Hauser, 2022). However, these neuromodulatory influences are expressed through distributed cortical circuits, suggesting that variability in the structural properties of such circuits controls how neurochemical signals are integrated and translated into behaviour. These findings highlight the potential for individual-specific neurobiological foundations of decision-making, yet further research is necessary to confirm this link between biology and behaviour.

Numerous previous studies have identified specific brain regions and functional connectivity pathways underlying key decision variables. In particular, the ventromedial prefrontal cortex (vmPFC) has been repeatedly implicated in choice probability, valuation, and subjective utility of available choices (Cockburn et al., 2022; Daw et al., 2006). The frontopolar cortex has also been highlighted in many studies as a locus of exploratory behaviours and choice switching (Boorman et al., 2009; Daw et al., 2006), and the posterior cingulate cortex (PCC) has been implicated in tracking value computations across diverse decision-making contexts (Clithero & Rangel, 2014). In addition to functional pathways, brain structural properties such as cortical thickness have been recently shown to contribute to decision-making processes (Filmer et al., 2023; Smid et al., 2023; Sunderaraman et al., 2022) and are subject to age-related changes (Liu et al., 2025). The behavioural variability we observe in decision-making contexts may therefore result from inter-individual differences in brain structure, which can arise from developmental or genetic sources, and in turn influence function (Frank et al., 2009). Examining such cortical differences through microstructural imaging can potentially reveal the underlying neuroanatomical origins of these observed differences in behaviour and brain anatomical pathways. More specifically, exploration-exploitation behaviours rely on the balance between flexible, stochastic policy control, often associated with prefrontal networks, and more stable, value-based action selection frequently attributed to parietal and sensorimotor regions (Daw et al., 2006; Klein-Flügge & Bestmann, 2012). Such processes are likely to rely heavily on the efficiency of neural signalling and underlying structural connectivity.

Quantitative MRI is a valuable method for the quantification and investigation of *in vivo* brain microstructures, providing a non-invasive means of assessing neuroanatomical features relevant to cognitive function (Callaghan et al., 2014; Weiskopf et al., 2015; Ziegler et al., 2019). These indices extend beyond conventional morphometric measures such as cortical volume or thickness to index variation in quantitative, non-arbitrary units with direct histological correlates. For example, previous work has exploited these recent advances in quantitative neuroimaging to correlate model parameters of respiratory interoception with *in vivo* indicators of cortical histology, revealing multiple neuroanatomical contributions to interoceptive sensitivity, precision, and metacognition (Nikolova et al., 2025). In particular, the R2* quantitative contrast, which indexes the iron content of cortical tissues, has been repeatedly associated with noradrenergic and dopaminergic activity due to the accumulation of the neuromelanin-iron complex in noradrenergic neurons in the locus coeruleus, and dopaminergic neurons in the substantia nigra (Riley et al., 2023; Zucca et al., 2017). Noradrenaline is broadly recognised as a key contributor to higher-level cognitive processes including decision-making and uncertainty processing (Hauser et al., 2017; Lawson et al., 2021; Yu & Dayan, 2005). Therefore, investigating this relationship between structural brain components and neurotransmitters with putative roles in decision-making may advance our understanding of why decision-making behaviours differ so greatly between individuals.

In this study, we aimed to examine the neuroanatomical foundations of inter-individual differences in exploration-exploitation behaviours. To this end, 122 participants completed a gamified exploration task, constructed as a multi-armed bandit, which enabled the quantification of subject-level variability in exploration-exploitation behaviours (Dubois et al., 2021; Dubois and Hauser, 2022). We applied a whole-brain, cluster-corrected multiple linear regression analysis on three distinct neuroanatomical maps generated through quantitative MRI, relating cortical architectures with task parameter estimates characterising exploration-exploitation behaviours. Through associating individual-specific model parameter estimates denoting behavioural heuristics with cortical microstructures, we sought to elucidate the neurobiological bases of decision-making processes, and how they might drive inter-individual differences in learning.

## Materials and Methods

### Participants

A total of 566 (360 females, 205 males, 1 other) participants (median age = 24, age range = 18 - 56) participated in the Visceral Mind Project, a large-scale neuroimaging project at the Center of Functionally Integrative Neuroscience, Aarhus University. Participants were recruited through local advertisements such as social media, posters, and flyers, and also via the SONA online participation pool system. Participants were required to have normal or corrected-to-normal vision and fluency in Danish or English. Exclusion criteria required that participants were not taking any medications excluding contraceptives or over-the-counter antihistamines. In addition, participants were required to be compatible with standard MRI scanning requirements (i.e., not pregnant/breastfeeding, no metal implants, no claustrophobia, etc.). Participants from this dataset completed multiple behavioural tasks, physiological recordings, and MRI scanning, in addition to psychiatric and lifestyle inventories, across three separate visits scheduled on different days. The MRI and behavioural data reported in this study were collected on different days. The study was granted ethical approval by the local Region Midtjylland Ethics Committee and was conducted in accordance with the Declaration of Helsinki (2013). All participants provided written informed consent and were remunerated for their participation.

### Exploration-Exploitation Paradigm – ‘*Maggie’s Farm*’

‘*Maggie’s Farm*’ is a multi-armed bandit paradigm designed to examine human exploration-exploitation strategies in decision-making (Dubois et al., 2021; Dubois & Hauser, 2022) (**Figure 1C**). On each trial, participants were asked to choose between three “bandits” (represented by trees bearing apples), each associated with a given reward distribution (apple size), to maximise their total reward. Each bandit’s reward (apple size) was drawn from a normal distribution with a fixed sampling variance. Bandits displayed either high prior reward information (i.e., three initial samples), or limited prior information (one initial sample), while one novel bandit provided no prior reward information. Bandits also had either a standard or a low reward mean. At the start of each trial, participants were shown initial reward samples displayed on a wooden crate at the bottom of the screen. They then selected a bandit to sample from. Participants were instructed to collect the biggest apples before the end of each trial (sunset). Participants could perform either one draw (short horizon) or six draws (long horizon) from the selected bandit, depending on the trial condition. By analysing participants’ choices, the task distinguishes between different exploration strategies, such as selecting novel options to reduce uncertainty or choosing randomly without regard for expected value. See Supplementary Information for detailed task instructions, and Dubois et al. (2021) for further information regarding details of the paradigm.

**Figure 1.**
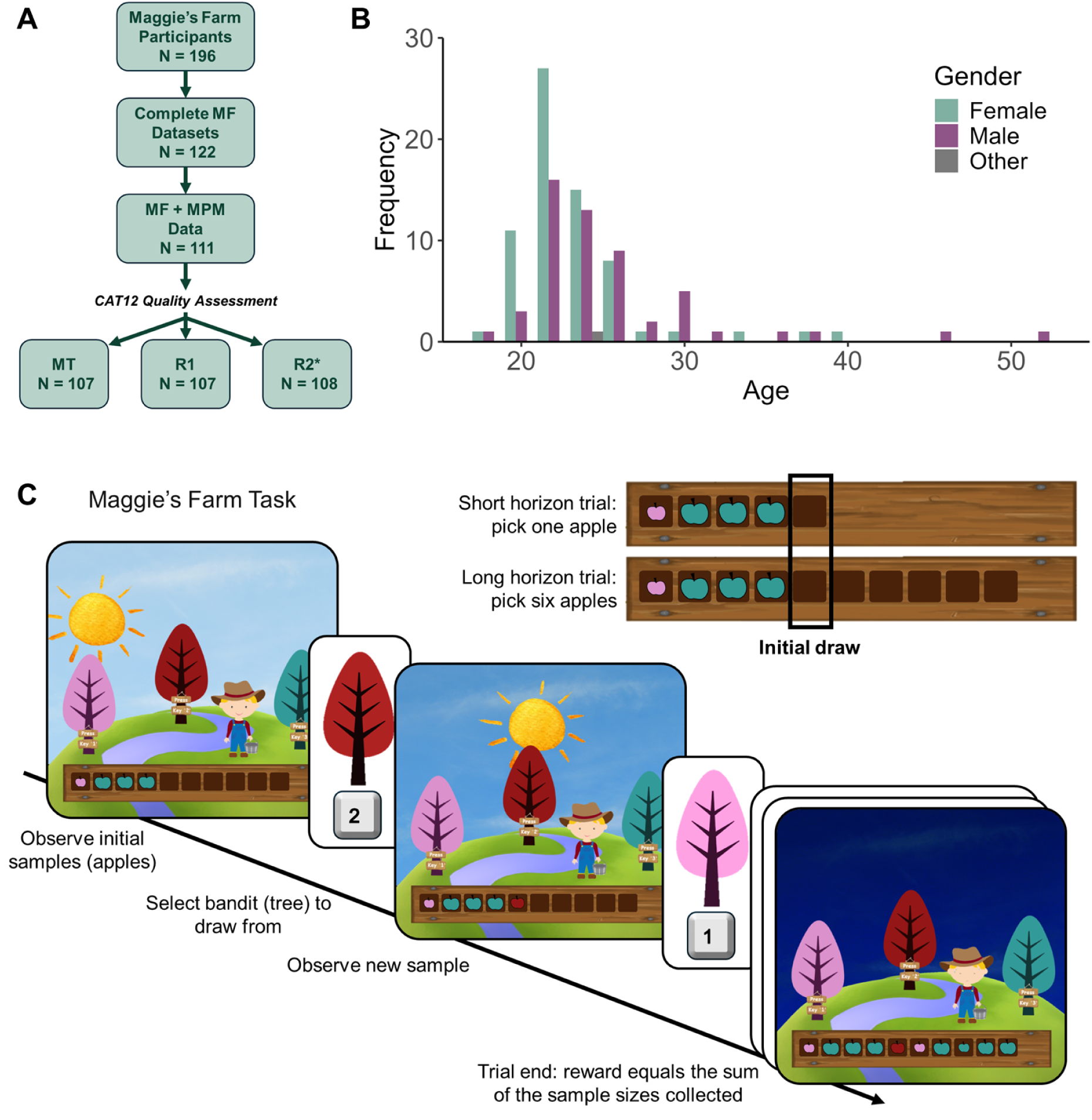
Participant inclusion and demographics. (**A**) Participant inclusion diagram, detailing data quality controls applied to both the behavioural and neuroimaging data. (**B**) Distributions of age and gender of the 122 participants included in the behavioural analyses. (**C**) Maggie’s Farm (MF) task design. Participants could choose from three possible bandits (trees) to obtain samples (apples) of varying value (apple size), to maximise their reward. At the start of each trial, participants were shown initial samples in the crate at the bottom of the screen. *Top right:* trial horizon and number of remaining draws were indicated by the number of empty crates. Our analyses primarily focused on the initial draw for both horizon conditions.

### Statistical Analyses of Behaviour

To first assess normality in participants’ choice behaviour, behavioural heuristics, and model parameter estimates, we conducted Shapiro-Wilk normality tests. Due to non-normal distributions in the behavioural data, we proceeded with non-parametric statistical analyses for all comparisons and correlations. To examine whether trial horizon manipulation influenced exploratory behaviours (and as positive controls to demonstrate consistency with the findings of Dubois et al. 2021; Dubois and Hauser 2022), we compared the choice frequencies of different bandit values in the long versus short horizon trials using two-sided Wilcoxon signed-rank two-tailed tests. To identify potential correlations between behavioural metrics and model parameters, we conducted partial Spearman’s rank correlation analyses correcting for age, gender, and body mass index (BMI) and correcting for multiple comparisons using false discovery rate (FDR) correction (see **Supplementary Figure 1** for partial correlation matrix). To investigate the association between participants’ choice behaviours and model heuristics, and mental health factors extracted from psychiatric surveys, we applied partial Spearman’s rank correlation analyses, correcting for age, gender, and BMI and correcting for multiple comparisons using FDR correction (see **Supplementary Information, Supplementary Figures 2 and 3** for results, factor structure, and further details).

### Computational Modelling of Behaviour

The computational model used here employs Thompson sampling (capturing uncertainty-driven value-based random exploration), with the addition of both value-free random exploration and novelty exploration following the framework described in Dubois et al. (see Dubois et al. 2021; Dubois and Hauser 2022, for further model details).

This previous validation work examining the same task in a different sample (Dubois et al., 2021) compared multiple computational models, consisting of three base models: the UCB model which includes the UCB algorithm and a softmax choice function; the Thompson model which employs Thompson sampling; and a hybrid model which combined the UCB model and the Thompson model. These computationally-demanding models were compared both with and without the addition of simpler heuristic exploration strategies, i.e., value-free random exploration (ε-greedy), and novelty exploration (novelty bonus η). Here, we employ the winning model from this model comparison, i.e., the model with the greatest held-out data likelihood (%): the Thompson model with the addition of ε-greedy and novelty bonus η.

To model value-free random exploration in line with these previous analyses, we added an ε-greedy component to the decision rule, ensuring that every ε% of the time, an option alternative to the one predicted is chosen. Also in line with previous work, we added a novelty bonus η to the computation of the value of novel bandits in order to model novelty exploration.

The value of each bandit, *i*, is defined as follows:

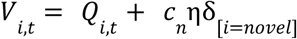

A sample 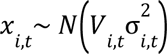 is taken from each bandit. The probability of choosing bandit *i* depends on the probability that all pairwise differences between the sample from bandit *i* and the other bandits *j*≠*i* were greater or equal to 0 (see Speekenbrink & Konstantinidis, (2015) for the probability of maximum utility choice rule). In *Maggie’s Farm*, two pairwise differences scores (contained in the two-dimensional vector *u*) were computed for each bandit due to the presence of three bandits at a time. The probability of choosing bandit *i* is described by:

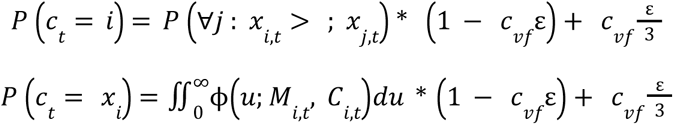

where ϕ is the multivariate Normal density function with mean vector:

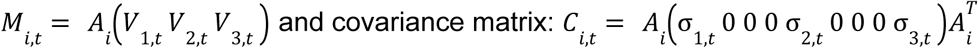

where the matrix *A_i_* computes the pairwise differences between bandit *i* and other bandits. For example, for bandit *i* = 1:

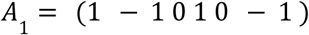

We used the *maximum a posteriori* (MAP) probability estimate for fitting parameter values, employing the fmincon optimisation function in MATLAB, in line with previous work (see Dubois et al. 2021; Dubois and Hauser 2022, for further details regarding parameter estimation).

### Multi-Parameter Brain Mapping

The multi-parameter pipeline employed here follows that described in Nikolova et al. (2025). We utilised a well-established qMRI protocol (Weiskopf et al., 2013, 2015) to map percent saturation resulting from magnetisation transfer (MT), longitudinal relaxation rate (R1) and effective transverse relaxation rate (R2*), followed by voxel-based quantification to examine how subject-specific parameter estimates of exploration-exploitation behaviours correlate with different characteristics of brain microstructure.

#### Data Acquisition

Neuroimaging data were acquired using a 3T MR system (Magnetom Prisma, Siemens healthcare, Erlangen, Germany), using a standard 32-channel radiofrequency (RF) head coil and a body coil. We obtained high-resolution whole brain T1-weighted anatomical images (0.8 mm^3^ isotropic) using an MP-RAGE sequence (repetition time = 2.2 s, echo time = 2.51 ms, matrix size = 256 x 256 x 192 voxels, flip angle = 8°, AP acquisition direction).

We acquired whole-brain image acquisitions at isotropic 0.8 mm resolution using an MPM quantitative imaging protocol (Callaghan et al., 2019; Weiskopf et al., 2013). The sequences comprised three spoiled multi-echo 3D fast low angle shot (FLASH) acquisitions and three additional calibration sequences to correct for RF transmit field inhomogeneities. The FLASH sequences were acquired with MT, PD and T1 weighting. We used a flip angle of 6° for the MT- and PD-weighted images, and 21° for the T1-weighted acquisitions. MT-weighting employed a Gaussian RF pulse 2 kHz off resonance with 4 ms duration and a nominal flip rate of 220°. The field of view was 256 mm head-foot, 224 mm anterior-posterior, and 179 mm right-left. Gradient echoes with alternating readout gradient polarity were obtained using equidistant echo times ranging from 2.34 to 13.8 ms (MT) or 18.4 ms (PD and T1), using a readout bandwidth of 490 Hz/pixel. For the MT-weighted acquisition, only six echoes were collected to achieve a repetition time (TR) of 25 ms for all FLASH volumes. For accelerated data acquisition, partially parallel imaging was performed using the GRAPPA algorithm, with an acceleration factor of 2 in each phase encoded direction and 40 integrated reference lines. A slab rotation of 30° was used for all acquisitions. The B1 mapping acquisition comprised 11 measurements with the nominal flip rate ranging from 115° to 65° in 5° steps. The total scanning time for the qMRI acquisitions was approximately 26 minutes.

#### Map Creation

All qMRI images were pre-processed using the hMRI toolbox v. 0.5.0 (January 2023) (Tabelow et al., 2019) and SMP12 (version 12.r7771, Wellcome Trust Centre for Neuroimaging, http://www.fil.ion.ucl.ac.uk/spm/), to correct the raw qMRI images for spatial transmit, receive field inhomogeneities and obtain quantitative MT, PD, R1 and R2* estimate maps. This correction was executed using empirical data, i.e., the RF sensitivity map and B1 maps collected from each participant. Excluding enabling imperfect spoiling correction, the hMRI toolbox was configured using default settings. All images were reoriented to MNI standard space prior to map creation. This processing produced four maps modelling different aspects of tissue microstructure: an MT map sensitive to myeloarchitectural integrity (Helms et al., 2008), a PD map representing tissue water content, an R1 map reflecting myelination, iron concentration and water content (primarily driven by myelination) (Lutti et al., 2014), and an R2* map sensitive to tissue iron concentration (Langkammer et al., 2010).

MT saturation and R1 both reflect qualities of myelin, however they index different underlying biophysical properties (Filo et al., 2019; Weiskopf et al., 2021). R1 is predominantly shaped by tissue composition, water content, and iron, whereas MT saturation indexes macromolecular content through magnetisation transfer. As a result, subtle variations in microstructural myelin components may exert a greater effect on one map type compared with the other. However, the in-depth understanding of the neuroarchitecture-to-MPM contrast association remains an active research area, and some biophysical constructs contribute to both map types.

The unified segmentation approach (Ashburner & Friston, 2005) was used to segment MT saturation maps into grey matter (GM), white matter (WM) and cerebrospinal fluid (CSF) probability maps. We used tissue probability maps based on multi-parametric maps developed by Lorio et al. (2016), without bias field correction given that MT maps do not show significant bias field modulation. We then used the GM and WM probability maps to perform inter-subject registration using Diffeomorphic Image Registration (DARTEL), a nonlinear diffeomorphic algorithm (Ashburner, 2007). The MT, PD, R1 and R2* maps were then normalised to MNI space (at isotropic 1 mm resolution) using the resulting DARTEL template and participant-specific deformation fields. The nonlinear registration of the quantitative maps was based on the MT maps due to their high contrast in subcortical structures, and a WM-GM contrast in the cortex similar to T1 weighted images (Helms et al., 2009). Finally, tissue-weighted smoothing was applied using a kernel of 8 mm full width at half maximum (FWHM) using the voxel-based quantification (VBQ) approach (Draganski et al., 2011). Importantly, in contrast to voxel-based morphometry analysis, this VBQ smoothing approach aims to minimise partial volume effects and optimally preserves the quantitative values of the original qMRI images by not modulating the parameter maps to account for volume changes. The resulting GM segments for each map were used for all statistical analyses.

### MRI Quality Control and Participant Exclusions

Multi-parameter mapping contrast images were acquired for 503 total participants. Following inspection, several participants were removed from all analyses for reasons related to either MRI or behavioural data. Three participants were excluded immediately following MR data collection for medical reasons (one cerebral palsy and two other suspected brain abnormalities). Three participants were excluded due to errors in the scanning sequences. As MPM data is known to be particularly sensitive to motion artefacts, we conducted a thorough quality control (QC) assessment to identify high-motion images. Visual QC HTML reports were created of each participant using the hMRI-vQC toolbox (Sherif et al., 2022), and all reports were visually inspected and labelled by two researchers. Doubtful cases were discussed, and a further 57 participants were excluded due to excessive motion affecting the tissue-class segmentation. Data from the remaining 443 participants was used in the spatial analysis and template creation using DARTEL.

A total of 196 participated in the *Maggie’s Farm* task, 122 of which had complete datasets for the task (median age: 24, age range: 18-52, 67 females, 54 males, 1 other). Participants with missing or incomplete behavioural data were excluded from these analyses. The overlap between the behavioural paradigm and MPM data left 111 participants for VBQ analyses. Following additional QC using the CAT12 toolbox in SPM12, a further four participants were excluded from VBQ analyses for MT maps (MT: final N = 107), four for R1 maps (R1: final N = 107), and three for R2* maps (R2*: final N = 108) (**Figure 1**).

### Voxel Based Quantification Analysis

Grey and white matter masks were generated based on our samples, by averaging the smoothed, modulated GM and WM segment images, and thresholding the result at *p* > .2. Inter-subject variation in MT, R1 and R2* GM maps were modelled in separate multiple linear regression analyses. A total of 16 regressors of interest were used in the VBQ analysis, consisting of behavioural heuristics and subject-specific model parameter estimates generated through Thompson sampling modelling (see **Table 2** for a full list of regressors included). Additionally, we included age, gender, body mass index (BMI) and total intracranial volume (TIV) as nuisance covariates in all analyses, following recommended procedures for computational neuroanatomy (Ridgway et al., 2008). VBQ data is often subject to non-stationarity, therefore we applied Threshold-Free Cluster Enhancement (TFCE) correction to our contrasts in combination with a GM mask, which provides non-parametric statistics and is robust to such non-stationarity in the data. We analysed whole-brain maps of each positive and negative *t*-contrast using a TFCE-corrected FWE-cluster *p*-value with *p* < .05 inclusion threshold (Hupé, 2015; Ridgway et al., 2008). All statistical analyses were conducted in SPM12, and we utilised the JuBrain Anatomy Toolbox v. 3.0 (Eickhoff et al., 2005) to determine anatomical labels and regional percentages.

**Table 1:** Participant demographics of 122 *Maggie’s Farm* participants.

| Variable | Mean (± SD) |
| --- | --- |
| Age | 24.9 ± 5.0 |
| Gender | 67 (54.9%) female, 54 (44.3%) male, 1 (0.8%) other |
| BMI | 23.1 ± 2.9 |

**Table 2:** Behavioural model parameter glossary.

| Parameter | Definition |
| --- | --- |
| $\epsilon_i$ SH | Epsilon greedy parameter for short horizon trials |
| $\epsilon_i$ LH | Epsilon greedy parameter for long horizon trials |
| $\eta$ SH | Novelty bonus for short horizon trials |
| $\eta$ LH | Novelty bonus for long horizon trials |
| $\sigma_0$ SH | Prior variance for short horizon trials |
| $\sigma_0$ LH | Prior variance for long horizon trials |
| $Q_0$ | Expected value before seeing any samples |
| Score SH | Score in the short horizon (i.e., the size of the single apple picked) |
| Score all LH trials | Total score in the long horizon (i.e., the sum of all the apples picked) |
| Score first LH trial | Score on the first trial in the long horizon (i.e., the size of the first apple picked) |
| Behaviour EV SH | Expected value of chosen bandit (i.e., exploitation), in the short horizon |
| Behaviour EV LH | Expected value of chosen bandit (i.e., exploitation), in the long horizon |
| Behaviour IS SH | Number of samples that were displayed for the bandit that they chose (i.e., information seeking), in the short horizon |
| Behaviour IS LH | Number of samples that were displayed for the bandit that they chose (i.e., information seeking), in the long horizon |
| Behaviour Consist SH | Consistency measure, i.e., whether participants made the same choice when observing the same data, in the short horizon |
| Behaviour Consist LH | Consistency measure, i.e., whether participants made the same choice when observing the same data, in the long horizon |

**Table 3:** Summary of whole brain VBQ results: negative correlation between MT Saturation and ε-greedy parameter in short horizon trials.

| Region | Significant Whole Brain Clusters |  |  |  |  |  |
| --- | --- | --- | --- | --- | --- | --- |
| | $k$ | $p(\text{FWE-corr})$ | TFCE | x | y | z |
| R Postcentral Gyrus | 5840 | 0.024 | 2681.63 | 68.8 | -10.4 | 20 |
| R Superior Parietal Lobule | 3883 | 0.036 | 2373.76 | 24.8 | -46.4 | 47.2 |
| L Cerebellum | 757 | 0.038 | 2340.27 | -10.4 | -59.2 | -36 |
| R Middle Frontal Gyrus | 1181 | 0.041 | 2289.43 | 53.6 | 14.4 | 40 |
| R Medial Orbital Gyrus | 620 | 0.044 | 2239.23 | 16 | 44 | -20 |
| R Anterior Orbital Gyrus | 528 | 0.045 | 2219.54 | 24 | 40.8 | -11.2 |
| R Supramarginal Gyrus | 404 | 0.047 | 2200.75 | 63.2 | -24.8 | 39.2 |
| L Superior Occipital Gyrus | 676 | 0.047 | 2193.52 | -16 | -82.4 | 37.6 |
| R Superior Frontal Gyrus | 97 | 0.049 | 2167.19 | 20.8 | 58.4 | 4.8 |
| R Middle Frontal Gyrus | 70 | 0.049 | 2157.75 | 48 | 44.8 | 17.6 |

**Table 4:** Summary of whole brain VBQ results: positive correlation between R1 and ε-greedy parameter in long horizon trials.

| Region | Significant Whole Brain Clusters |  |  |  |  |  |
| --- | --- | --- | --- | --- | --- | --- |
| | $k$ | $p(\text{FWE-corr})$ | TFCE | x | y | z |
| R Superior Frontal Gyrus | 6539 | 0.015 | 3083.99 | 8 | -11.2 | 77.6 |
| R Middle Frontal Gyrus | 3126 | 0.023 | 2742.65 | 36 | 12.8 | 55.2 |
| R Precentral Gyrus | 7143 | 0.025 | 2683.08 | 39.2 | -8.8 | 60.8 |
| R Cerebellum | 14419 | 0.026 | 2662.38 | 8 | -68.8 | -30.4 |
| L Postcentral Gyrus | 2334 | 0.030 | 2542.83 | -47.2 | -24 | 59.2 |
| L Precentral Gyrus | 3629 | 0.031 | 2520.42 | -8.8 | -16.8 | 69.6 |
| L Middle Frontal Gyrus | 1698 | 0.033 | 2477.97 | -31.2 | 13.6 | 50.4 |
| R Middle Temporal Gyrus | 1150 | 0.045 | 2253.25 | 50.4 | -48 | 2.4 |
| R Middle Frontal Gyrus | 653 | 0.046 | 2236.41 | 48.8 | 13.6 | 42.4 |
| R Angular Gyrus | 496 | 0.047 | 2217.32 | 46.4 | -65.6 | 33.6 |
| L Middle Frontal Gyrus | 347 | 0.047 | 2210.99 | -39.2 | 43.2 | -2.4 |
| L Middle Frontal Gyrus | 580 | 0.048 | 2202.08 | -34.4 | 39.2 | 18.4 |

## Results

### Exploration-Exploitation Behaviours are Modulated by Trial Horizon

To evaluate participants’ chosen exploration-exploitation strategies, we first analysed participants’ choice behaviour in the multi-armed bandit task, *Maggie’s Farm*, and estimated subject-specific behavioural heuristics through Thompson sampling modelling (**Figure 1C**).

Participants chose to sample less from the high-value bandit in long horizon trials compared with short horizon trials (two-sided Wilcoxon signed-rank two-tailed test: *V* = 336.5, *p* < .001), demonstrating that participants were willing to forego selecting the bandit with the greatest expected reward outcome to potentially learn about alternative bandits (**Figure 2A**). Consequently, participants chose to sample more from the novel bandit (*V* = 514.0, *p* < .001) and from the low-value bandit (*V* = 1076.5, *p* < .001) in long horizon trials versus short horizon trials (**Figure 2A**). This demonstrates that participants chose to engage in more exploratory behaviours in the long horizon trials, where exploration is beneficial, compared with short horizon trials, where exploration is less beneficial. This behaviour is also consistent with that presented in previous studies examining choice behaviours in the same task (Dubois et al., 2021; Dubois & Hauser, 2022), which found that manipulating trial horizon drove exploratory behaviours.

**Figure 2.**
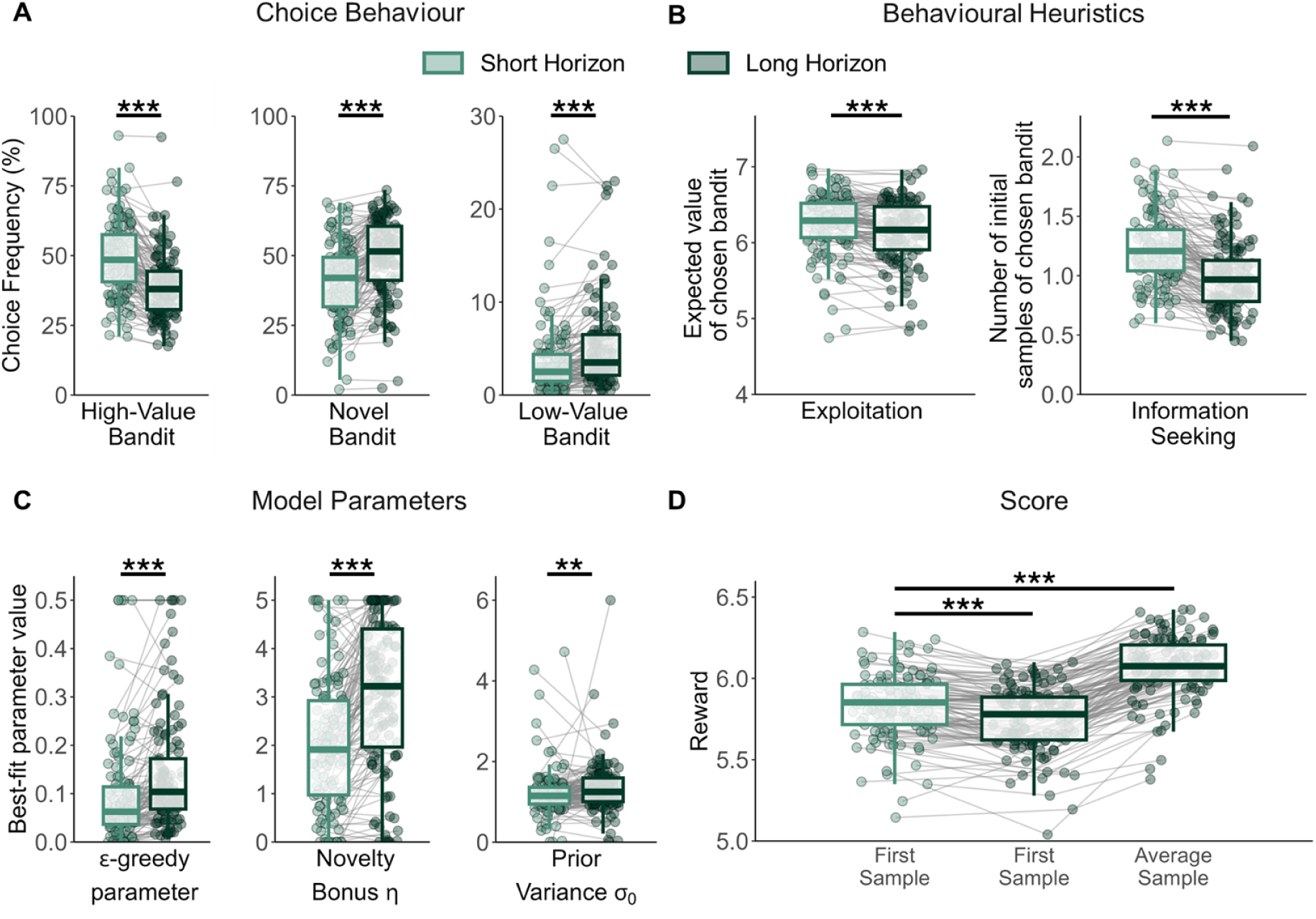
Effects of trial horizon on bandit choice behaviour (**A**), behavioural heuristics (**B**), Thompson sampling model parameters (**C**), and reward earned in the first sample and average sample within a trial (**D**). **A:** In long horizon trials, participants sampled significantly less from the high-value bandit (*left*), and more from the novel bandit (*centre*) and low-value bandit (*right*). **B:** *Exploitation:* participants selected bandits with lower expected values more in the long versus short horizon, i.e., participants explored more in the long horizon. *Information-seeking:* participants chose lesser-known bandits more frequently in the long versus short horizon. **C:** In long horizon trials compared with short horizon trials, participants had significantly higher estimates of ε (value-free random exploration; *left*), η (novelty bonus; *centre*), and σ_0_ (prior variance; *right*). Parameters were fitted separately for each horizon and were fitted to the first draw of each participant. **D:** Participants obtained lower rewards in the initial draw of long horizon trials compared to short horizon trials, suggesting that participants initially prioritised acquiring information over rewards. This information gathering resulted in higher rewards overall in long horizon trials, as the newly acquired information enabled participants to make more informed decisions on bandit selection. **\*\*\*** indicates *p* < .001, **\*\*** indicates *p* < .01.

To investigate this choice behaviour further, we generated behavioural metrics describing exploitation behaviours (expected value of the selected bandit, i.e., mean value of the initial displayed samples) and information seeking behaviours (i.e., number of samples displayed for the selected bandit). If participants exhibited more explorative behaviours, we would expect participants to initially select bandits with lower expected values in long horizon trials. As expected, we found that participants chose bandits with a lower expected value (i.e., they explored more) in the long horizon trials compared with the short horizon trials (*V* = 5640.0, *p* < .001) (**Figure 2B**). Participants also chose lesser known (i.e., more informative) bandits more frequently in the long horizon trials versus short horizon trials (*V* = 6861.0, *p* < .001) (**Figure 2B**), suggesting that participants were more likely to select bandits they were less familiar with in the long horizon trials, potentially to acquire information and resolve uncertainty about unfamiliar bandits.

### Participants utilise value-free exploration when exploration may be beneficial

To formally quantify how participants employ different exploration strategies, we examined differences in behavioural model parameter estimates between long and short horizon trials. The ε-greedy parameter indexes value-free random exploration, in that ε% of the time each available bandit has an equal probability of being selected by the participant. Participants had higher values of ε-greedy, i.e., greater value-free random exploration, in the long horizon versus short horizon trials (two-sided Wilcoxon signed-rank two-tailed test: *V* = 1167.0, *p* < .001) (**Figure 2C**), suggesting that participants employed value-free explorative strategies in trials where exploration was beneficial. Participants also had significantly greater novelty bonuses, i.e., they showed greater novelty exploration in long horizon trials versus short horizon (*V* = 643.0, *p* < .001) (**Figure 2C**). Additionally, participants exhibited higher prior variances, i.e., uncertainty-driven value-based random exploration, or greater uncertainty about a bandit’s mean before seeing any samples, in long horizon trials versus short horizon trials (*V* = 2624.0, *p* = 0.00398) (**Figure 2C**), further bolstering the suggestion that exploration strategies were promoted during long horizon trials, which allowed the participant to subsequently exploit the information gathered to obtain greater rewards overall. These findings also remain consistent with those in Dubois et al. (2021, 2022), further demonstrating that long horizon trials engendered robust value-free exploratory behaviours.

### Value-free random exploration leads to greater rewards in the long run

To establish whether this choice behaviour yielded greater rewards in the long run, we examined reward earned in the initial samples of short horizon trials, initial samples in long horizon trials, and average reward earned across all six samples in the long horizon. As expected, participants earned fewer rewards in the first sample of long horizon trials versus short horizon trials due to the horizon-specific choice behaviour described above (FDR-corrected two-sided Wilcoxon signed-rank two-tailed test: *V* = 1015.5, *p* < .001) (**Figure 2D**). However, participants earned higher rewards on average across all samples in the long horizon compared to the short horizon initial sample (*V* = 10.0, *p* < .001), suggesting that participants initially selected less optimal bandits to gather information about lesser-known bandits, then applied this newly acquired information to select more optimal bandits in the future and maximise their rewards.

### Voxel-Based Quantification reveals neuroanatomical correlates of value-free exploration

We next investigated how these behavioural metrics and model parameter estimates of value-free random exploration in our exploration-exploitation paradigm relates to brain microstructural indices. To achieve this, we employed whole-brain multiple linear regression with TFCE correction against participants’ ε-greedy parameter estimates in short horizon trials and long horizon trials separately, while adjusting for age, gender, BMI, and total intracranial volume (TIV).

We found that R1 map values in the right superior frontal gyrus (TFCE corrected: *k* = 15138, *p*FWE_corr_ = .013, peak voxel coordinates: x = 8.0, y = -11.2, z = 77.6) and right middle frontal gyrus (TFCE corrected: *k* = 3467, *p*FWE_corr_ = .021, peak voxel coordinates: x = 36.0, y = 12.8, z = 55.2) were significantly positively correlated with ε-greedy parameter estimates specifically in long horizon trials (**Figure 3A-C**). In contrast, we found no significant correlations between R1 map values and ε-greedy values in short horizon trials. This finding specifically in long horizon but not short horizon trials therefore indicates a significant positive association between value-free random exploration in trials where exploration is more beneficial, and myelination in right frontal gyri, which have been previously linked to the control of impulsive behaviours (Hu et al., 2016).

**Figure 3.**
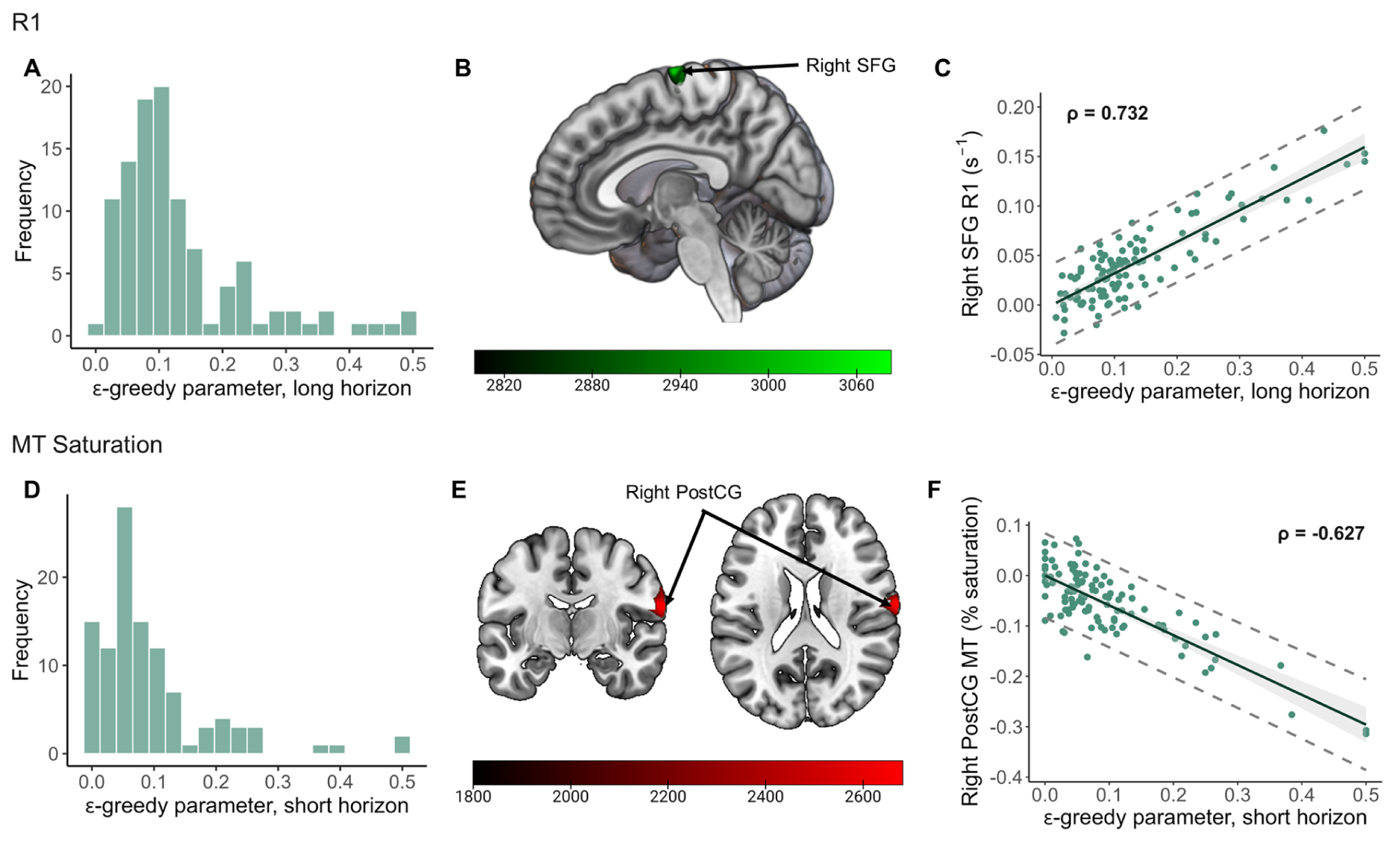
Microstructural correlates of value-free random exploration (ε-greedy) for R1 (**A**-**C**) and MT maps (**D**-**F**). **A:** Distribution of ε-greedy parameter estimates in long horizon trials across the cohort (N = 107). **B-C:** R1 values in the right superior frontal gyrus (SFG) are positively correlated with ε-greedy estimates in the long horizon. **D:** Distribution of ε-greedy parameter estimates in short horizon trials across the cohort (N = 107). **E-F:** MT saturation in the right postcentral gyrus (PostCG) is negatively correlated with ε-greedy estimates in the short horizon. Maps were TFCE (non-parametric) cluster-corrected for data non-stationarity and FWE corrected for multiple comparisons (*p*FWE < .05). Colour bars indicate TFCE values, scatter plots are for visualisation purposes.

We also found that MT saturation in the right postcentral gyrus (TFCE corrected: *k* = 5840, *p*FWE_corr_ = .024, peak voxel coordinates: x = 68.8, y = -10.4, z = 20) showed significant negative correlation with ε-greedy estimates in the short horizon trials (**Figure 3D-F**). MT saturation in the right superior parietal lobule was also negatively correlated with ε-greedy estimates in short horizon trials (TFCE corrected: *k* = 3883, *p*FWE_corr_ = .036, peak voxel coordinates: x = 24.8, y = -46.4, z = 47.2). However, we found no significant associations between MT saturation and ε-greedy estimates in long horizon trials. In short horizon trials, participants make only one bandit choice, therefore the optimal bandit choice in the short horizon is to select the bandit with the greatest expected value and to forego value-free random exploration and information seeking. These findings therefore indicate that greater myeloarchitectural integrity in the right postcentral gyrus and right superior parietal lobe is associated with reduced value-free random exploration when such exploration is not beneficial.

These significant clusters were only detectable in association with ε-greedy parameter estimates and were not detectable when examining bandit selection frequencies (see Supplementary Figure 4, Supplementary Tables 1-2), suggesting that our computational model captures distinct neurobiological effects compared with those captured by simpler behavioural metrics, such as raw selection frequencies.

No significant associations between behavioural heuristics of value-free random exploration and R2* map values survived TFCE correction, suggesting no obvious relationship between exploration-exploitation behaviours and cortical iron concentration.

### Meta-analytic Coactivation analyses uncover functional associations

Finally, we conducted a meta-analytic coactivation analysis of our findings in the right postcentral gyrus for MT map values and right SFG for R1 map values. To do this we utilised Neurosynth; a publicly-available meta-analytic database, using the right SFG and right postcentral gyrus peaks as seed regions. We generated meta-analytic maps displaying significant co-activation with our seed regions, in addition to associated functional and psychological terms from the literature. We sorted all associated terms with a non-zero reverse inference z-score by their posterior probability of appearing in a publication, given a pattern of activation (p(term | activation)). The top literature terms (max. ten) for each region are presented in Supplementary Tables 3-4 (see Supplementary Figure 6 for meta-analytic maps for each region).

We found that the right postcentral gyrus displayed significant coactivation with the right primary visual cortex, and bilateral premotor regions. In terms of functional associations, the right postcentral gyrus was most commonly related to speech, in line with suggestions that the right postcentral gyrus is linked to sensory representations of the larynx (example top terms: “pitch”, “speech production”; “vocal”) (Belyk & Brown, 2014). Previous work has also implicated functional activations in the right postcentral gyrus in implicit learning in autism spectrum disorder (Schipul & Just, 2016). In contrast, the right SFG displayed significant coactivation with the right insula, in addition to left premotor/supplementary motor regions. The right SFG was primarily associated with motor-based meta-analytic terms (example top terms: “muscle”, “motor premotor”), and has been linked to error detection and inhibition of distractor interference (Chevrier & Schachar, 2010; Xu et al., 2011).

## Discussion

Here, we utilised the *Maggie’s Farm* gamified multi-armed bandit task to interrogate the microstructural neural correlates of value-free random exploration behaviours. We applied Thompson sampling with additional behavioural heuristics describing value-free random exploration and novelty (Krebs et al., 2009) to examine how participants selected different behavioural strategies. We found that participants specifically made use of long horizon trials to sample lesser-known bandits and acquire new information to increase their overall reward, closely replicating previous findings which apply these same behavioural heuristics to the same task (Dubois et al., 2021; Dubois & Hauser, 2022). Further, we employed whole-brain regression analyses to demonstrate significant associations between heuristics of value-free exploration in long horizon trials and indices of myeloarchitecture in right superior frontal areas, which have been previously implicated in impulsive behaviours (Hu et al., 2016; Lim et al., 2021; P. Zhang et al., 2023). Specifically, we utilised voxel-based quantification (VBQ), which enables the biophysical, voxel-wise analysis of quantitative brain maps denoting indicators of iron concentration and myelin content (Draganski et al., 2011). In contrast to voxel-based morphometry (VBM), which supplies indirect, relative differences in tissue densities and is sensitive to differences in processing choices, VBQ provides biophysically plausible parameters describing absolute microstructural variations in cortical tissues (Senjem et al., 2005). Together, our findings here suggest that individual differences in decision-making and exploration behaviours may have distinct microstructural foundations in the brain.

Here, we employ a computational model to quantitatively describe specific psychological traits such as value-free random exploration, by combining the resource-demanding Thompson sampling with two computationally-light heuristic strategies. The utilisation of multiple different exploration behaviours, which all approximate an optimal exploration strategy, has been widely reported in human decision-making (Dubois et al., 2022; Gershman, 2018; Wilson et al., 2014). Examining simpler behavioural metrics such as bandit selection frequencies does not necessarily allow us to tease apart these different properties of exploration and may absorb shared variance from multiple forms of exploratory behaviour, for example directed versus random exploration.

Previous work has robustly demonstrated that participants employ both complex strategies and simpler heuristic strategies to seek and gather information during trials in which they can subsequently exploit this information (Dubois et al., 2021; Dubois & Hauser, 2022; Wilson et al., 2014). In this study, we successfully replicated these findings by combining simple heuristic modelling of value-free random exploration (ε-greedy) and novelty bonuses (η) with the more complex and resource-demanding Thompson sampling model. We found that participants chose low-value bandits significantly more in long horizon trials compared to short horizon trials, demonstrating that participants prioritised acquiring information about low-value or alternative bandits rather than maximising their reward in the short term by selecting high-value bandits in the first instance. We also found that participants chose lesser-known bandits more frequently in long horizon versus short horizon trials, indicating that participants took advantage of the long horizon to gather information about more informative (lesser-known) bandits to potentially maximise their reward in later samples, in line with previous studies (Warren et al., 2017; Wilson et al., 2014; Wu et al., 2018). We confirmed this hypothesis by examining participants’ mean scores for their initial samples in both the long and short horizons, and the average score across all six samples in the long horizon. In line with our hypotheses and with previous findings (Dubois et al., 2021; Dubois & Hauser, 2022), participants earned significantly more rewards across long horizon trial samples compared with short horizon initial samples, even though participants earned significantly less reward in long horizon initial samples.

Through whole-brain regression analyses, we then demonstrated positive associations between cortical myelination in the right superior (SFG) and right middle frontal gyri (MFG), including the supplementary motor area (SMA), and value-free random exploration in long horizon trials only. Since the SMA lies within the superior frontal gyrus, our region of significant correlation may reflect contributions from both regions. The right SFG has been implicated in impulsivity (Hu et al., 2016; P. Zhang et al., 2023), whereas the SMA is more classically associated with action planning and language processing (Hartwigsen et al., 2013; Hertrich et al., 2016; Moore-Parks et al., 2010). Our findings presented here may therefore depict an interaction between cognitive control and motor planning processes.

The link between increased value-free exploration and increased myelination in the SMA has not been previously reported. However, increased activation in the pre-SMA and striatum have been shown to facilitate faster decision-making by lowering response thresholds and alleviating global inhibition from the motor system (Forstmann et al., 2008). Further, sequential sampling models have demonstrated that activation in the pre-SMA, caudate nucleus, and anterior insula play important roles in evidence accumulation over time, which in turn facilitates value-based decision-making (Gluth et al., 2012). Therefore, these findings point to potential contributions of greater microarchitectural integrity in the SMA to action planning and evidence accumulation in time-pressured decision-making contexts. However, future studies are required to more closely examine the specific inputs of the SMA to these cognitive processes.

Previous work has identified greater activations in the right SFG during exploratory behaviours compared with exploitative behaviours in a classic four-armed bandit task (Laureiro-Martínez et al., 2014). A recent systematic review highlighted contributions of bilateral superior and middle frontal gyri to exploration-exploitation behaviours: specifically, the right and left middle frontal gyri were identified as core regions contributing to greater exploratory behaviours compared to exploitation, with right and left superior frontal gyri as secondary regions (Wyatt et al., 2024). Further, recent research examining brain structure has shown that cortical thickness in the SFG is negatively associated with impulsivity, indicating that impulsiveness, which is commonly associated with various psychiatric disorders, may rise from neuroanatomical differences (Lim et al., 2021; Schilling et al., 2013). Together, this highlights a putative role for myelination in right frontal regions in exploration behaviours. However, future studies specifically probing the link between cortical microstructures and impulsive behaviours, possibly combining impulsivity-focused behavioural tasks with impulsivity self-report scales, would provide vital additional information to confirm this link.

We also showed that value-free exploration in short horizon trials was negatively correlated with indices of cortical myeloarchitecture in the right postcentral gyrus, in that lower magnetisation transfer (MT) saturation values were associated with greater value-free random exploration and vice versa, specifically in short horizon trials in which this exploration is not necessarily beneficial. We observed this negative effect in the right postcentral gyrus; a region of the primary somatosensory cortex postulated to exhibit sensory representations of the larynx (Belyk & Brown, 2014). This association between exploratory behaviours and brain microstructures in the right postcentral gyrus is a novel finding which has not previously been documented. Recent work has, however, presented evidence for strong positive correlations between postcentral gyrus activity and the value of reducing variability, indicating that the postcentral gyrus may play a role in how an agent values the resolution of uncertainty (S. Zhang et al., 2025). In combination with our findings here, this may point towards a role for macromolecular content in the primary somatosensory cortex in uncertainty reduction. Additionally, a recent VBM study demonstrated a negative association between grey matter volume in the postcentral gyrus and perceptual sensitivity of numerosity (Yuan et al., 2023). Previous work has also found that the structure of primary sensory brain regions plays a crucial role in visual acuity and sensitivity on an individual level (Duncan & Boynton, 2003; Schwarzkopf et al., 2011). These findings, along with our findings in this study, suggest that structural variability in primary sensory brain areas may contribute to inter-individual differences in conscious experience, which may, in turn, beget variability in how individuals process information in their observed environment to inform their decision making.

The association between variations in cortical microstructures and exploration-exploitation behaviours may stem from various mechanisms. Myelination, for instance, is responsible for increased conduction velocity of axonal electrical signals, and is therefore crucial for facilitating fast and efficient neural signalling (Nave, 2010; Purves et al., 2001). Speculatively, this improved signalling efficiency may enable more flexibility in individuals’ internal models of the environment, therefore facilitating switching to exploratory behavioural strategies in response to environmental instability. The right SFG has been previously implicated with exploratory decision-making (Laureiro-Martínez et al., 2014; Wyatt et al., 2024), whereas the right postcentral gyrus has been linked to sensory gating and regulation of task-relevant sensorimotor responses (Meehan & Staines, 2007; Staines et al., 2002). Within this framework, increased myelination in the right SFG may indicate enhanced efficiency in the neural circuits supporting exploratory policy implementation, consistent with higher ε-greedy parameter estimates in trials where exploration is beneficial. In contrast, greater indices of myeloarchitecture in the right postcentral gyrus may improve the suppression of task-irrelevant sensorimotor signals and the regulation of stochastic response tendencies, thereby reducing unnecessary random exploration and supporting more stable exploitative behaviour in trials where exploration is not beneficial.

Interestingly, our analyses did not reveal any significant correlations between value-free exploration and canonical dopaminergic midbrain or hippocampal regions, despite their well-established roles in exploration-exploitation behaviours and decision-making (Rigoli et al., 2016; Schwartenbeck et al., 2015). The absence of significant associations in these regions may be a consequence of our rigorous whole-brain multiple regression pipeline with correction for multiple comparisons, which may lead to reduced sensitivity to subtler effects in smaller brain regions of interest. Moreover, it has been recently reported that qMRI parameters vary across cortical depths and correlate with layer-specific gene expression and cell counts (McColgan et al., 2021). To further investigate the contributions of subcortical microstructural variation to exploration behaviours, future studies may employ optimised qMRI sequences to specifically target subcortical regions of interest (Carter et al., 2025; Chen et al., 2014). For example, multiple-measurement protocols have been previously employed to improve the low signal-to-noise ratio typically linked to the high spatial resolution necessary to image smaller subcortical structures, such as the locus coeruleus and substantia nigra (Chen et al., 2014).

A limitation of this study is the correlational design in which behavioural and neural data were acquired separately, therefore one cannot make inferences about potential causal pathways linking brain structure and function to behavioural metrics and psychiatric symptom scores. Future research could involve identifying genetic contributions to psychiatric symptoms and differences in brain structures, and examining how such polygenic influences may impact various aspects of decision-making. Further, longitudinal studies investigating how lifestyle, environment, and mental health may impact inter-individual variability in cortical microstructures may provide valuable insights into the complex interplay between brain structure, brain function, and exhibited behaviours. Despite recruiting participants displaying a range of psychiatric phenotypes, we did not explicitly study clinical populations. Previous work has shown brain microstructural changes in Parkinson’s disease patients (Drori et al., 2025), and differences in white matter microstructures in cognitively healthy individuals who express strong genetic factors of late-onset Alzheimer’s disease (Operto et al., 2018). A natural next step from the work presented here is to examine how decision-making strategies in individuals suffering from mental health conditions that impact decision-making such as ADHD, depression, and anxiety, may be reflected in microstructural signatures in the brain.

As with any voxel-based methodology, our findings may be subject to some degree of sensitivity to processing choices, including spatial normalisation and smoothing parameters. To mitigate this, we followed established preprocessing guidelines used in other previous VBQ studies (Nikolova et al., 2025; Vejloe et al., 2026). Specifically, we employed a unified segmentation-normalisation framework in which registration and tissue classification are performed simultaneously (Ashburner & Friston, 2005), and utilised a smoothing approach specifically optimised for VBQ to minimise partial volume effects and optimally preserve the quantitative values of the original qMRI images (Draganski et al., 2011).

The variation in qMRI parameters across cortical depths presents an additional limitation, as the processing pipeline employed here does not account for depth-specific effects. Future work could build upon our voxel-level analyses by utilising surface-based methods, such as surface-based registration, to examine such depth-specific effects and also improve the precision of spatial normalisation (Khan et al., 2011).

Previous research has indicated that exploration model parameters may lack generalisability (Eckstein et al., 2022; Witte et al., 2025), which presents a possible limitation to the current study. However, the exploration parameters in our model do not denote general exploration, rather they index specific exploratory behaviours; ε-greedy for example specifically denotes value-free random exploration, and the novelty bonus η indexes novelty exploration. Also, our previous study (Dubois & Hauser, 2022) employing the same computational model and task in a different sample also collected self-report survey scores in the Barratt Impulsiveness Scale (BIS), and Adult ADHD Self-Report Scale (ASRS). This work demonstrated significant positive correlations between ε-greedy parameter estimates in long horizon trials and general impulsivity, motor impulsivity (subscale of BIS), ADHD, and ADHD hyperactivity-impulsivity (subscale of ASRS) scores (Dubois & Hauser, 2022). This builds upon previous findings linking exploration to increased impulsivity, particularly in ADHD (Hauser et al., 2014; Williams & Taylor, 2006), by demonstrating that value-free random exploration specifically is increased in ADHD, while associations were not found for other forms of exploration. This finding is congruent with the idea of impulsivity as ‘acting on a whim’, as value-free random exploration disregards all existing information and is therefore the least computationally-demanding. Future work may examine the association between variations in cortical microstructures and self-report measures of impulsivity, and how these metrics interact with various exploratory strategies. Additionally, we focused primarily on which bandit participants selected in their initial choice within a trial. Future work may also investigate at which point within long horizon trials participants switched from exploratory (i.e., selecting lower-value but novel or informative bandits) to exploitatory strategies to probe how cortical microstructures may drive individual differences in strategy switching.

Demographic factors such as socioeconomic status have been previously associated with exploration (Decker et al., 2025) and may therefore play an important role in individual differences in exploration-exploitation behaviours. The absence of such demographic data presents a potential limitation for this study, however our sample was primarily young adults recruited from the local University student population which in itself has relatively low socioeconomic variability compared to a community sample. Future work may employ exploratory factor analyses linking exploration behaviours and a range of lifestyle and demographic factors, including socioeconomic status.

Overall, we present novel insights into how behavioural heuristics of exploration-exploitation behaviours and decision-making are associated with distinct indices of cortical microstructures in task-relevant brain regions. Our findings here indicate that brain microstructures such as myelination may influence how individuals make decisions and act upon them to acquire information and seek rewards. This therefore highlights great potential for future work examining causal relationships between cortical microstructure, functional brain pathways, and observed behaviour, and may lay the groundwork for future studies examining how such cortical microstructures differ between individuals suffering from psychiatric conditions characterised by aberrant decision-making, such as anxiety disorders and ADHD.

## Acknowledgements

This research is financially supported by a Lundbeckfonden Fellowship (R272-2017-4345) and a European Research Council Grant (ERC-2020-StG-948788) awarded to MGA. The funding sources were not involved in the study design, collection, analysis, interpretation, or writing of the manuscript. TUH has received funding from the Wellcome Trust (316955/Z/24/Z), the European Research Council (ERC) under the European Union’s Horizon 2020 research and innovation programme (grant agreement No 946055), and the Carl-Zeiss-Stiftung. This work was supported by the Alexander von Humboldt foundation, (more) precisely the Alexander-von-Humboldt-Professorship award to Peter Dayan. For the purpose of Open Access, the author has applied a CC BY public copyright license to any Author Accepted Manuscript version arising from this submission.

## Supplementary Information

**Supplementary Figure 1.**
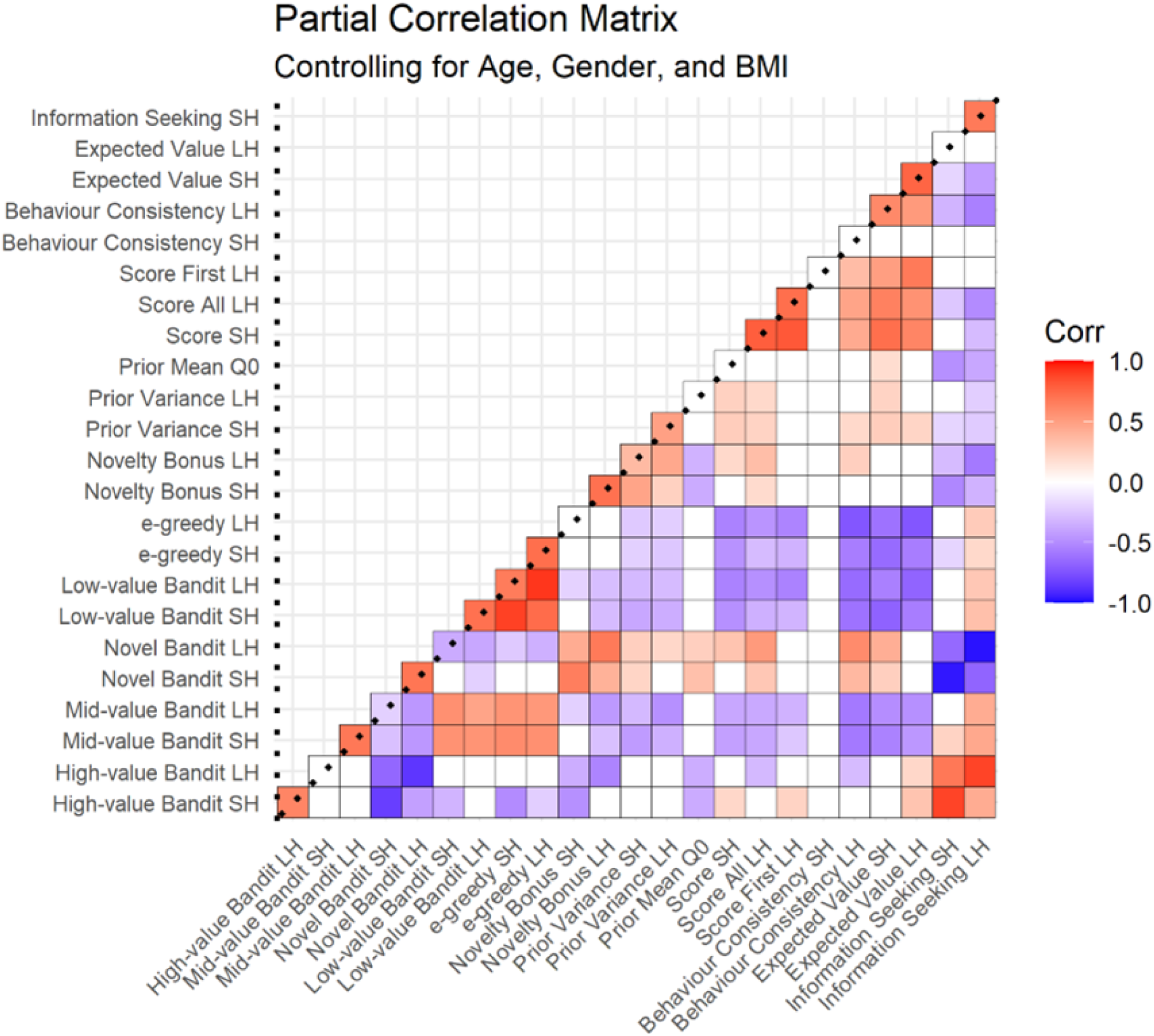
**SF1** Partial correlation matrix detailing Spearman’s correlations between choice behaviour, model heuristics, and Thompson model parameter estimates, corrected for age, gender, and body mass index (BMI).

### Supplementary Information 1: Associations between behavioural metrics and transdiagnostic psychiatric factors

To further interrogate the origins of participants’ exploration-exploitation behaviours, we examined the association between participants’ choice behaviours and model heuristics, and multi-level mental health factors. Factor scores for each participant denoting mental health constructs were drawn from item-level exploratory factor analyses (EFA) performed on responses from eight mental health self-report surveys from the full Visceral Mind Project cohort of 566 participants. Survey instruments spanned autism spectrum (AQ10), ADHD (ASRS), depression (MDI and PHQ9), somatic symptoms (PHQ15), stress (PSS), social anxiety (SIAS), and trait anxiety (STAI trait) (Banellis et al., 2025). This multi-level factor solution revealed 11 lower-level factors and two higher-level factors. The higher-level factor ‘ADHD & Somatic’ encompassed four lower-level factors: negative thoughts, restlessness, somatic symptoms, and impulsivity. The second higher-level factor ‘affective symptoms’ encompassed six lower-level factors: social anxiety, self-confidence, sleep, depression, stress, and autism (**Supplementary Figure 2**). The eleventh lower-level factor was inattentiveness, which we did not examine in this study (see Banellis et al. 2025 for further information regarding EFA methods).

We then applied Spearman’s rank partial correlations comparing behavioural metrics to mental health factors, correcting for age, gender, and body mass index (BMI), and correcting for multiple comparisons (FDR correction method, Benjamini-Hochberg procedure at < .05). We found a significant positive correlation between the affective symptoms higher-level factor and participants’ prior mean Q_0_, i.e., the prior beliefs held by a participant about a bandit’s mean value before seeing any samples (**Supplementary Figure 3**). This suggests that participants who exhibited greater affective symptoms of anxiety and depression were more likely to hold beliefs that unexplored bandits would yield higher rewards. This finding differs from previous work by Dubois and Hauser (2022) who instead found significant positive associations between the anxious-depression factor and participants’ novelty bonuses (Dubois and Hauser 2022).

We then conducted post-hoc Spearman’s rank partial correlations to further examine the relationship between the prior mean Q_0_ and the six lower-level factors encompassed within the affective symptoms higher-level factor, again correcting for age, gender, and BMI, and correcting for multiple comparisons (FDR correction). This revealed a significant positive correlation between the prior mean Q_0_ and the social anxiety lower-level factor, in addition to a significant negative correlation between the prior mean and the self confidence factor (negative loading) (**Supplementary Figure 3**). Together, these findings indicate that participants who experience greater social anxiety and reduced confidence in oneself may place inflated value expectations upon unknown options.

### Supplementary Information 2: Detailed Task Instructions

Prior to starting the task, participants underwent a short task tutorial with the following instructions, alongside visual examples of bandits (trees bearing apples) and rewards (apples):

“*Apples come in different shades and sizes. You need to help us collect the BIGGEST ones before sunset. You can only pick apples until sunset. Sometimes you will start at noon and will pick six apples, sometimes you will start in the afternoon and will pick only one apple. On each day you will collect apples from new trees. Some of the trees are better than others. To help you, some apples were already picked up before you arrived. These are the three different types of trees. Each tree produces an apple with a corresponding colour, but it’s their size that matters! For example, if we look at the apples produced, the yellow tree seems to be better on this day because it produces bigger apples ON AVERAGE. This is a big apple from the yellow tree. This is a medium apple from the red tree. This is a small apple from the yellow tree. This is a small apple from the blue tree. And finally this is a big apple from the blue tree. You will press the key 1, 2, or 3 to select the tree you wish to pick an apple from. If the sun is here, it is still early and you will be able to pick six apples. If the sun is almost set, you will only be able to pick one apple. At the end of each day, you will see the amount of juice extracted from the apples you collected. The juice will be poured in a small glass if one apple was picked, and in a big glass if six apples were picked.*”

**Supplementary Figure 2.**
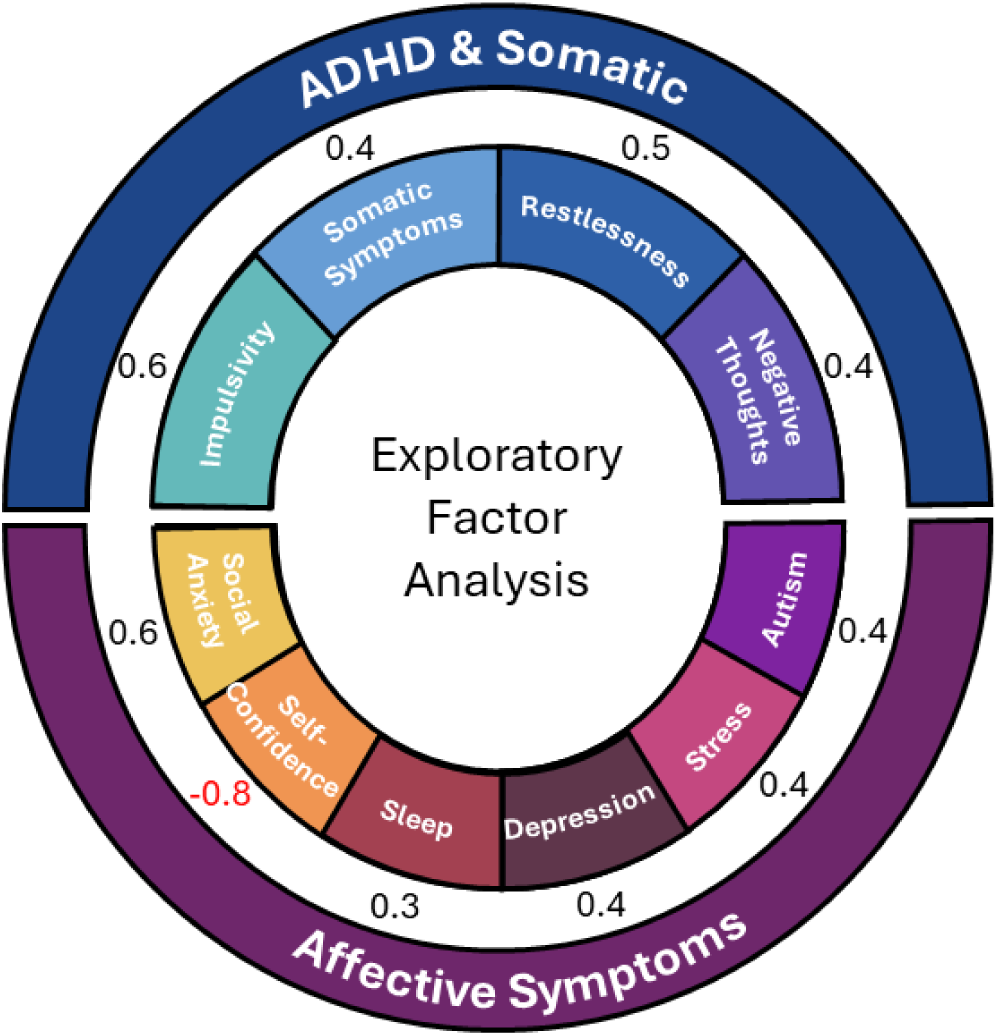
**SF2** Multi-level factor solution derived from the responses in eight mental health surveys of 566 participants in the Visceral Mind Project (Banellis et al. 2025). Outer circle denotes the two higher-level factors: affective symptoms and ADHD & somatic. Inner circle denotes the lower-level factors: social anxiety, self-confidence, sleep, depression, stress, autism, negative thoughts, restlessness, somatic symptoms, and impulsivity. The eleventh lower-level factor, inattentiveness, was not encompassed within either of the two higher-level factors and was not examined here. Numbers denote factor loadings for each lower-level factor. For more details regarding the exploratory factor analysis (EFA) methods, see Banellis et al. (2025).

**Supplementary Figure 3.**
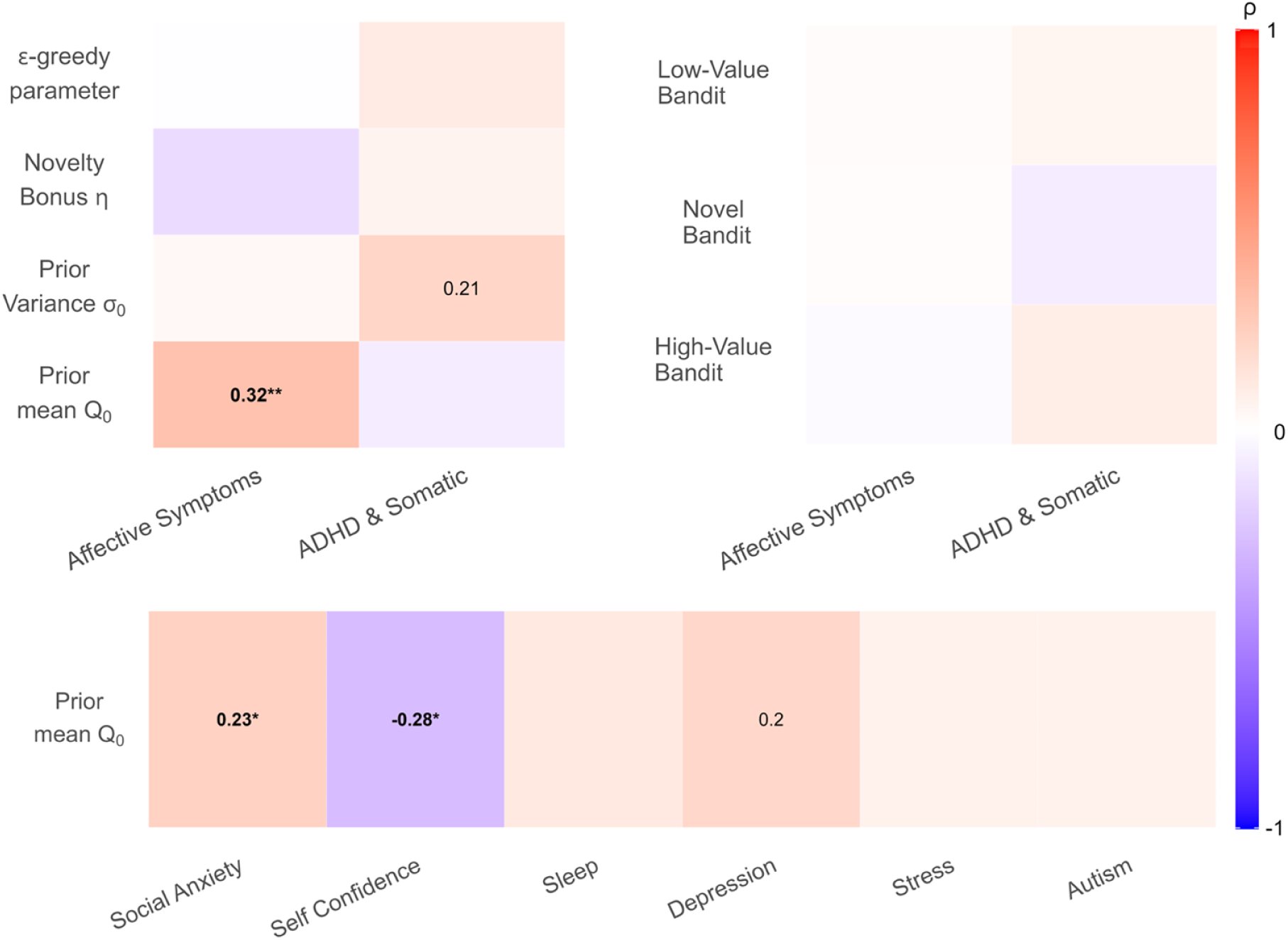
**SF3** *Left:* The exploratory factor analysis (EFA)-derived ‘Affective Symptoms’ higher-order factor was significantly correlated with prior mean Q_0_, i.e., participants’ prior beliefs about a bandit’s mean value before seeing any samples (r(107) = 0.32, *p_unc_* < .001, *p_cor_* = .00667; FDR-corrected partial correlations correcting for age, gender, and BMI). *Right:* There were no significant correlations between either of the EFA higher-order factors and participants’ behaviour (i.e., the frequency of picking the low-value, novel, or high-value bandits). *Bottom:* FDR-corrected post-hoc partial correlations correcting for age, gender, and BMI revealed that the association between the affective symptoms higher-order factor and prior mean Q_0_ is driven by significant positive correlation with the social anxiety factor (r(107) = 0.23, *p_unc_* = .0105, *p_cor_* = .0315) and significant negative correlation with the self confidence factor (r(107) = -0.28, *p_unc_* = .00248, *p_cor_* = .0149). For significant uncorrected correlations, *p* values are printed on the heatmaps, and *p* values for significant corrected correlations are indicated in bold with asterisks (\**p* < .05, \*\**p* < .01, \*\*\**p* < .001).

**Supplementary Figure 4.**
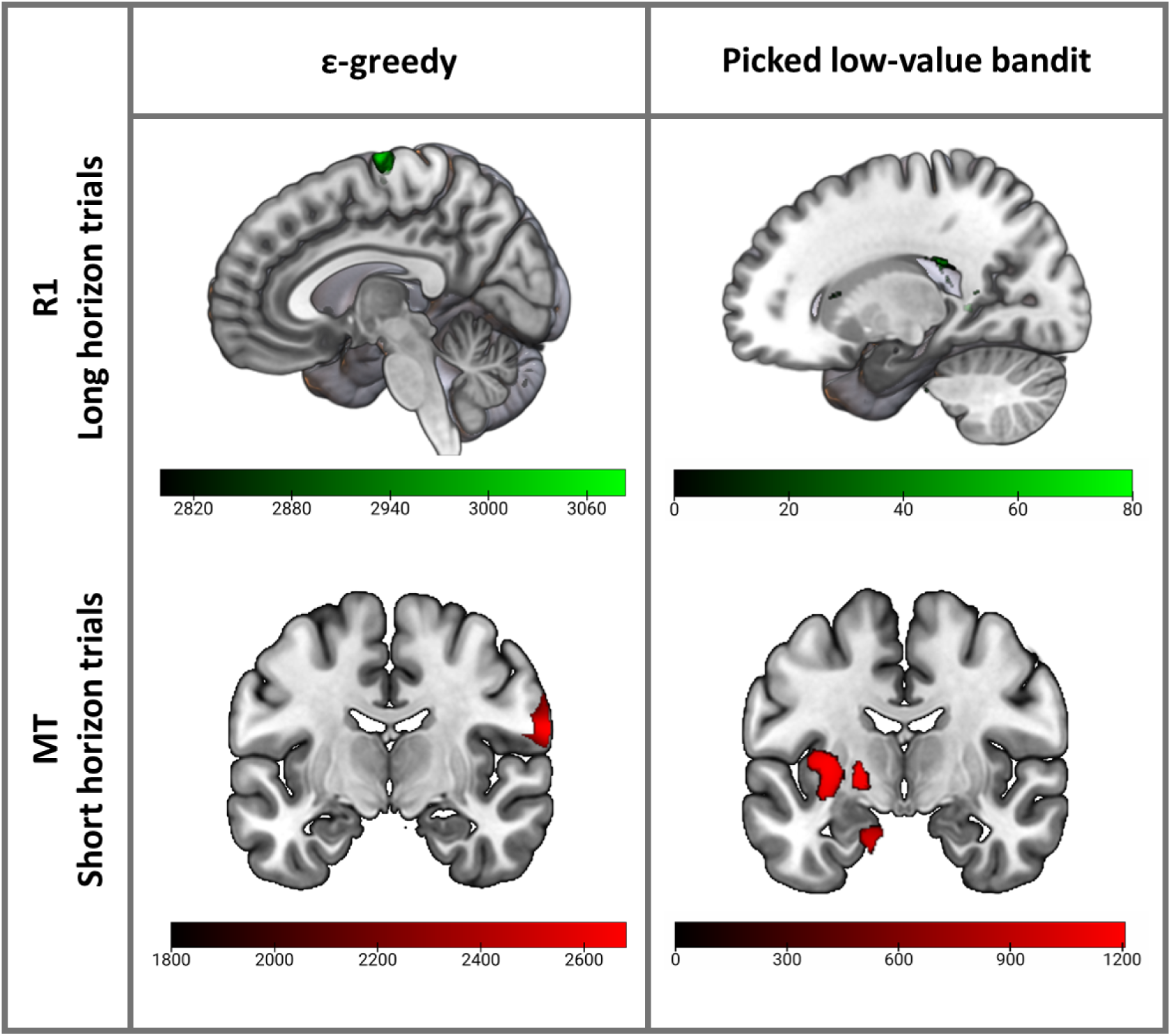
**SF4** Comparison of main VBQ findings examining associations between cortical microstructures and ε-greedy parameter estimates (left), versus the simpler behavioural metric of bandit selection frequency (right).

**Supplementary Figure 5.**
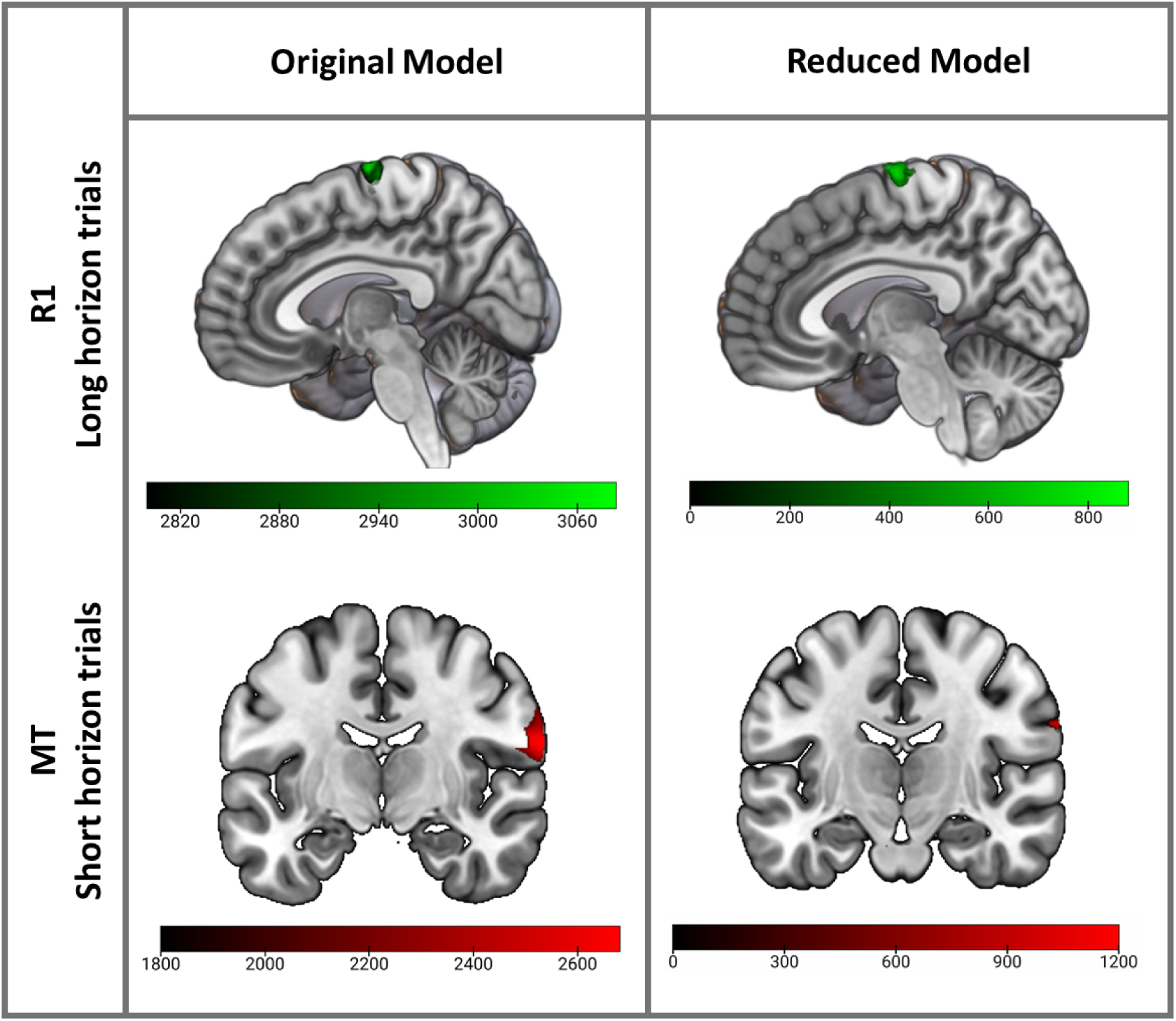
**SF5** Comparison of main VBQ findings for the original model comprising 16 regressors of interest and four nuisance covariates (age, gender, BMI, and TIV), and a reduced model comprising ε-greedy for short and long horizon trials only, plus the same four nuisance covariates.

**Supplementary Figure 6.**
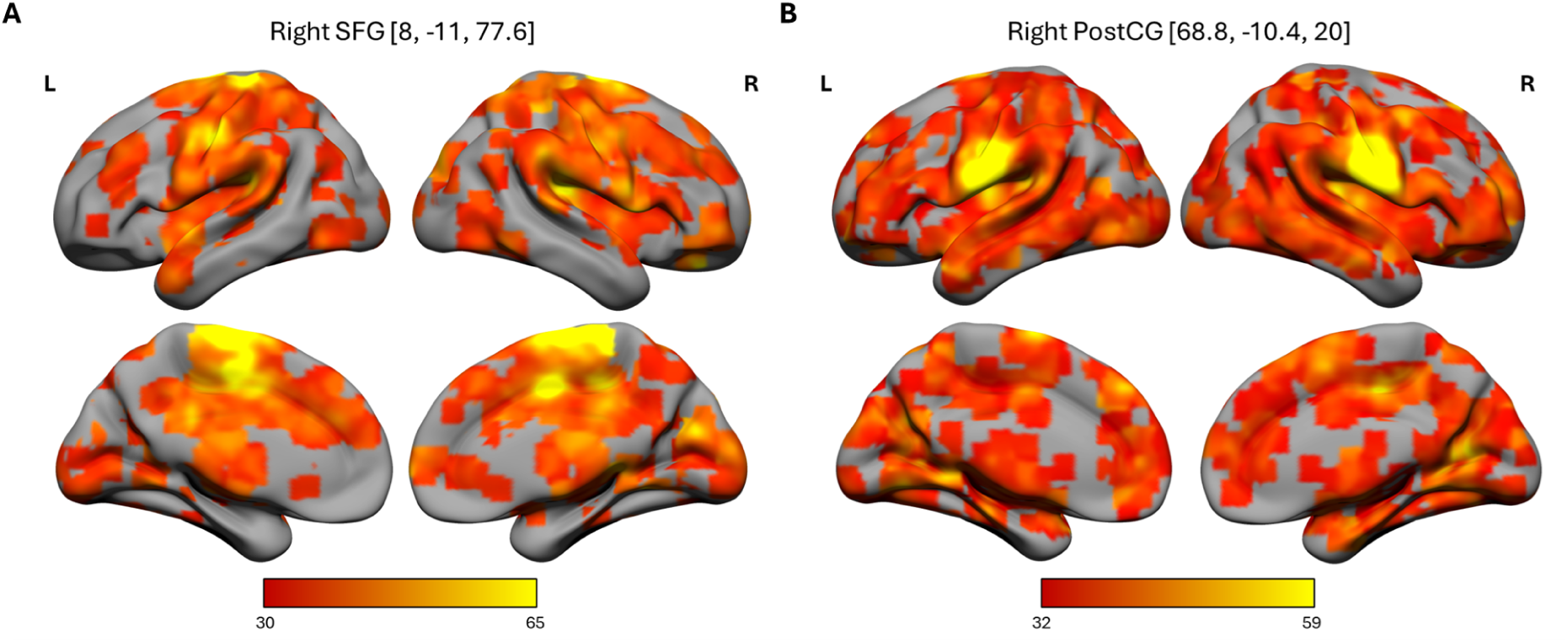
**SF6** Meta-analytic analyses for right SFG and right PostCG seed regions identified through VBQ analyses, displaying areas of significant coactivation. SFG = superior frontal gyrus; PostCG = postcentral gyrus.

## Supplementary Tables

**Supplementary Table 1:** Summary of whole brain VBQ results: positive correlation between R1 map values and low-value bandit selection frequency in long horizon trials. Clusters are ordered by FWE-corrected p value.

| R1 Whole Brain Clusters for Raw Behavioural Data |  |  |  |  |  |  |  |
| --- | --- | --- | --- | --- | --- | --- | --- |
| Region | k | p(FWE-corr) | p(uncorr.) | TFCE | x | y | z |
| Left lateral ventricle | 1 | 1.00 | 0.049 | 132.53 | -24.0 | -52.8 | 13.6 |
| Left lateral ventricle | 1 | 1.00 | 0.049 | 132.04 | -23.2 | -51.2 | 12.8 |
| Right lateral ventricle | 19 | 1.00 | 0.043 | 123.99 | 34.4 | -42.4 | 3.2 |
| Right lateral ventricle | 82 | 1.00 | 0.030 | 81.92 | 23.2 | -30.4 | 24 |

**Supplementary Table 2:**
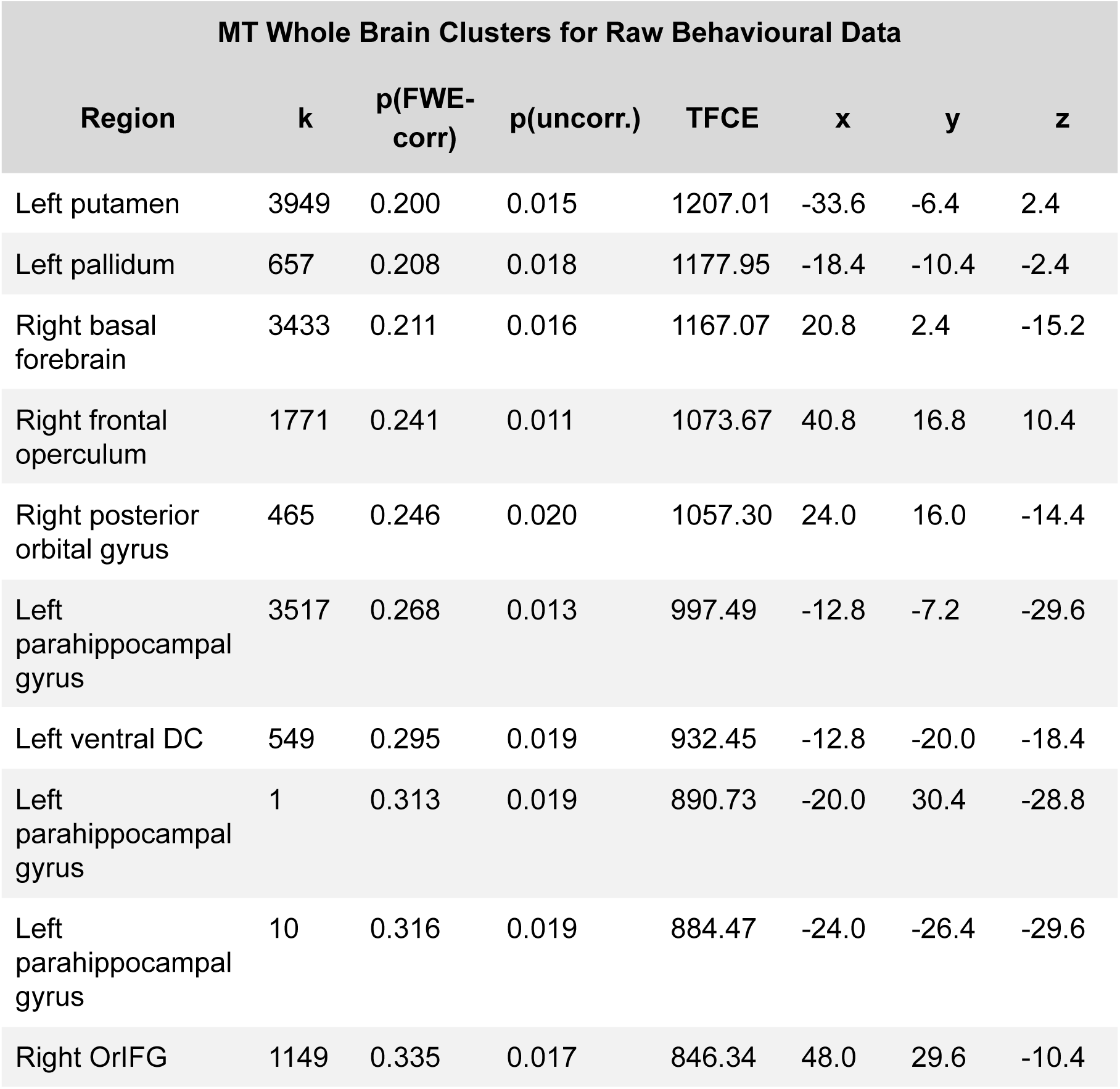
Summary of top ten whole brain VBQ results: negative correlation between MT Saturation and low-value bandit selection frequency in short horizon trials. Clusters are ordered by FWE-corrected p value. DC = diencephalon; OrIFG = orbital part of inferior frontal gyrus.

| MT Whole Brain Clusters for Raw Behavioural Data |  |  |  |  |  |  |  |
| --- | --- | --- | --- | --- | --- | --- | --- |
| Region | k | p(FWE-corr) | p(uncorr.) | TFCE | x | y | z |
| Left putamen | 3949 | 0.200 | 0.015 | 1207.01 | -33.6 | -6.4 | 2.4 |
| Left pallidum | 657 | 0.208 | 0.018 | 1177.95 | -18.4 | -10.4 | -2.4 |
| Right basal forebrain | 3433 | 0.211 | 0.016 | 1167.07 | 20.8 | 2.4 | -15.2 |
| Right frontal operculum | 1771 | 0.241 | 0.011 | 1073.67 | 40.8 | 16.8 | 10.4 |
| Right posterior orbital gyrus | 465 | 0.246 | 0.020 | 1057.30 | 24.0 | 16.0 | -14.4 |
| Left parahippocampal gyrus | 3517 | 0.268 | 0.013 | 997.49 | -12.8 | -7.2 | -29.6 |
| Left ventral DC | 549 | 0.295 | 0.019 | 932.45 | -12.8 | -20.0 | -18.4 |
| Left parahippocampal gyrus | 1 | 0.313 | 0.019 | 890.73 | -20.0 | 30.4 | -28.8 |
| Left parahippocampal gyrus | 10 | 0.316 | 0.019 | 884.47 | -24.0 | -26.4 | -29.6 |
| Right OrIFG | 1149 | 0.335 | 0.017 | 846.34 | 48.0 | 29.6 | -10.4 |

**Supplementary Table 3:** Top ten meta-analysis map associations for the right SFG, ordered by posterior probability.

| Name | z-score | Posterior prob. | Func. conn. (r) | Meta-analytic coact. (r) |
| --- | --- | --- | --- | --- |
| foot | 8.35 | 0.95 | 0.48 | 0.19 |
| muscle | 6.02 | 0.92 | 0.23 | 0.13 |
| suppressed | 5.95 | 0.92 | 0.01 | 0 |
| substantia | 5.13 | 0.92 | -0.05 | -0.01 |
| motor premotor | 4.82 | 0.91 | 0.28 | 0.12 |
| distractor | 4.75 | 0.91 | -0.01 | -0.01 |
| electrical | 5.12 | 0.9 | 0.19 | 0.16 |
| cortex dorsal | 4.58 | 0.9 | 0.02 | 0.01 |
| midbrain | 4.9 | 0.88 | -0.06 | 0 |
| coordination | 4.55 | 0.88 | 0.27 | 0.22 |

**Supplementary Table 4:**
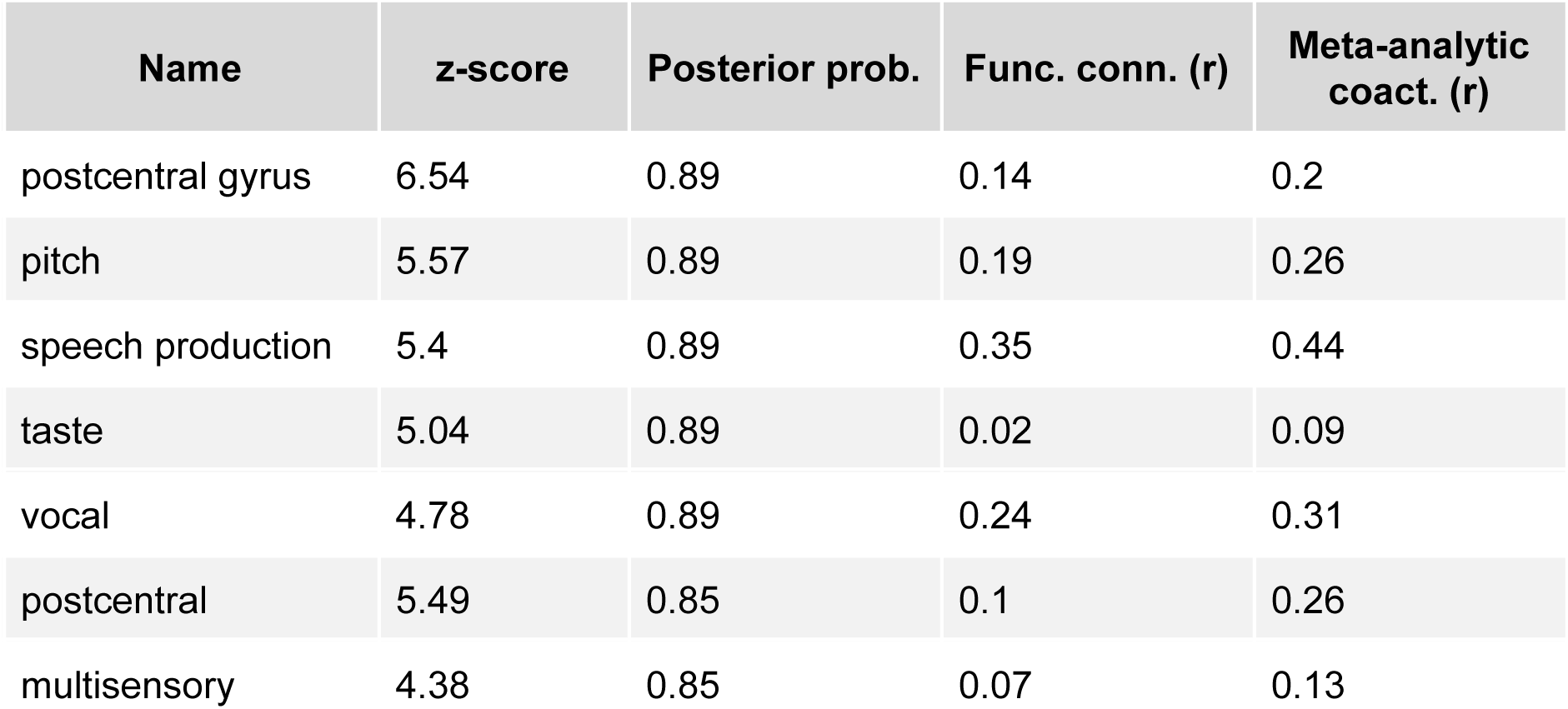
Meta-analysis map associations for the right postcentral gyrus, ordered by posterior probability.

